# Adjacent Neuronal Fascicle Guides Motoneuron 24 Dendritic Branching and Axonal Routing Decisions through Dscam1 Signaling

**DOI:** 10.1101/2024.04.08.588591

**Authors:** Kathy Clara Bui, Daichi Kamiyama

## Abstract

The formation and precise positioning of axons and dendrites are crucial for the development of neural circuits. Although juxtracrine signaling via cell-cell contact is known to influence these processes, the specific structures and mechanisms regulating neuronal process positioning within the central nervous system (CNS) remain to be fully identified. Our study investigates motoneuron 24 (MN24) in the *Drosophila* embryonic CNS, which is characterized by a complex yet stereotyped axon projection pattern, known as ‘axonal routing.’ In this motoneuron, the primary dendritic branches project laterally toward the midline, specifically emerging at the sites where axons turn. We observed that Scp2-positive neurons contribute to the lateral fascicle structure in the ventral nerve cord (VNC) near MN24 dendrites. Notably, the knockout of the Down syndrome cell adhesion molecule (*dscam1*) results in the loss of dendrites and disruption of proper axonal routing in MN24, while not affecting the formation of the fascicle structure. Through cell-type specific knockdown and rescue experiments of dscam1, we have determined that the interaction between MN24 and Scp2-positive fascicle, mediated by Dscam1, promotes the development of both dendrites and axonal routing. Our findings demonstrate that the holistic configuration of neuronal structures, such as axons and dendrites, within single motoneurons can be governed by local contact with the adjacent neuron fascicle, a novel reference structure for neural circuitry wiring.

**Significance Summary:** We uncover a key neuronal structure serving as a guiding reference for neural circuitry within the *Drosophila* embryonic CNS, highlighting the essential role of an adjacent axonal fascicle in precisely coordinating axon and dendrite positioning in motoneuron 24 (MN24). Our investigation of cell-cell interactions between motoneurons and adjacent axonal fascicles— crucial for initiating dendrite formation, soma migration, and axonal pathfinding in MN24— emphasizes the neuronal fascicle’s significance in neural circuit formation through Dscam1-mediated inter-neuronal communication. This enhances our understanding of the molecular underpinnings of motoneuron morphogenesis in *Drosophila*. Given the occurrence of analogous axon fascicle formations within the vertebrate spinal cord, such structures may play a conserved role in the morphogenesis of motoneurons via Dscam1 across phyla.

## Introduction

The positioning of axons and dendrites is crucial for neuronal function (Yogev and Shen, 2014; Lefebvre et al., 2015; Yogev and Shen, 2017; Lanoue and Cooper, 2019), particularly during early embryonic stages when neural circuits are formed independently of neuronal activities, under the guidance of the developmental program (Haverkamp, 1986; Verhage et al., 2000; Varoqueaux et al., 2002; Constance et al., 2018). Extracellular cues play a critical role in neural circuit development by providing spatial information that guides the growth and patterning of neural structures (Sanes, 1989; Guan and Rao, 2003; Long and Huttner, 2019). A well-studied example of spatial regulation in neuronal processes is seen in the decision-making regarding repulsion and attraction in reference to the midline within the embryonic central nervous system (CNS) of *Drosophila* melanogaster (Klambt et al., 1991; Evans and Bashaw, 2010; Howard et al., 2019). In the CNS, neurons can project their axons across the midline to the contralateral side or stay on the ipsilateral side (Kaprielian et al., 2001). This critical decision relies on the diffusible midline ligands like Slit and Netrin binding with their respective receptors Roundabout (Robo) and Frazzled (Fra) expressed on axons (Harris et al., 1996; Kidd et al., 1998; Brose et al., 1999; Kidd et al., 1999; Hiramoto et al., 2000). Similarly, these signaling mechanisms also direct the later, higher-order branches of dendrites in the CNS (Furrer et al., 2003; Furrer et al., 2007; Mauss et al., 2009). However, it remains to be determined whether these diffusible molecules from the midline exclusively regulate how neuronal processes are directed to their proper destinations within the embryonic CNS.

Current research identifies only a few adhesion molecules crucial for juxtracrine signaling in this context (Howard et al., 2019). For example, the atypical cadherin Flamingo is involved in axon midline crossing (Organisti et al., 2015), while Down syndrome cell adhesion molecule (Dscam1) promotes axon growth across segments (Alavi et al., 2016). Yet, their roles in CNS dendrite formation are largely unexplored. Our previous study fills this gap by demonstrating Dscam1’s significant role in dendritic outgrowth (Kamiyama et al., 2015). We investigated anterior corner cell (aCC) motoneurons, focusing on their lateral axonal extensions and interactions with MP1 partner neurons. We discovered that dendritogenesis is initiated by a cell-cell adhesion mechanism facilitated by Dscam1 interactions, where Dscam1 on one neuron binds to another, leading to cytoskeletal changes and dendritic growth in aCC motoneuron. The extent to which this Dscam1-mediated mechanism is generalizable among motoneurons, and its broader impact on Drosophila neuronal morphology, warrants further exploration. Additionally, evidence suggests Dscam1’s involvement in processes like cell body migration, axon guidance, and dendrite patterning, underscoring its broad involvement in neural development (Schmucker et al., 2000; Wojtowicz et al., 2004; Zhan et al., 2004; Zhu et al., 2006; Goyal et al., 2019; Liu et al., 2020; Dong et al., 2022; Wilhelm et al., 2022).

In the embryonic CNS, 36 motoneurons per hemisegment have been mapped (Sink and Whitington, 1991; Landgraf et al., 1997). These motoneurons are categorized into two groups: the first with cell bodies situated between the midline and the neuropiles along the mediolateral plane, and the second with cell bodies located outside the neuropiles, extending to the edge of the ventral nerve cord (VNC). The aCC motoneuron falls into this former category. In our current investigation, we shift our focus to the latter category to further explore Dscam1-mediated neural morphogenesis. Motoneurons in this second category exhibit an axon turning pattern and elaborate their dendrites specifically at these axonal turning points. One such motoneuron, MN24, displays a unique pattern of dendritic elaboration. Contrary to the aCC motoneuron, whose dendritic arbors are in the middle region of the neuropile, MN24’s dendritic projections are predominantly observed at the most lateral edge of the neuropile, without overlapping aCC processes. This positioning makes MN24 an ideal model for investigating the molecular and cellular mechanisms underlying axon turning and dendritic arborization in identifiable single motoneurons within the CNS.

We have anatomically characterized MN24 and its potential partner, a Scp2-positive neuronal fascicle. Our studies demonstrate that a dscam1 knockout eliminates dendrites and disrupts MN24 axonal routing. Cell-specific *dscam1* manipulations reveal that Dscam1-mediated contact between MN24 and Scp2-positive neurons is essential for both dendrite and axon development. These findings introduce a novel fascicle structure in which neuronal morphology is modulated through Dscam1 adhesion.

## Results

### Spatial Regulation of MN24 Dendritogenesis in Late Embryonic CNS

Previous studies have demonstrated that the dendritic processes of the MN24 are distinctly positioned away from those in the aCC motoneuron, exhibiting a notable lateral shift (Landgraf et al., 1997; Landgraf et al., 2003b). While these studies have characterized the dendritic morphologies of MN24 qualitatively, there are no current reports that carefully examine the exact positioning and detailed arrangements of its dendritic branches. In our prior study (Kamiyama et al., 2015), we identified a MN24-specific GAL4 driver, *hedgehog (hh)-GAL4*, during a screening of GAL4 drivers. Notably, this *hh-GAL4* driver initiates GAL4 expression at 11:00 AEL in MN24, as well as in several neighboring motoneurons (MN21, MN22, and MN23). Initially, using this GAL4 driver, we attempted to label neuronal processes by expressing a membrane marker (*UAS-mCD4::tdGFP*). However, due to the low expression level of this GAL4 driver, we were unable to achieve adequate membrane labeling to distinguish fine neuronal structures. Consequently, we decided to employ a retrograde lipophilic-dye labeling technique. This method allowed us to label membranes with a high density of lipophilic dye, enabling the detailed visualization of individual dendritic branches (Figure 1A). We then quantified the number and position of dendritic tips, the latter defined as the distance from the midline to the ventral nerve cord edge (**Figure 1A-B**). These measurements reveal that on average, wild-type MN24 at 15:00 AEL has 7.6 ± 0.3 (mean ± SEM) primary dendritic branches, which are located 15.8 ± 0.3 μm from the midline (**Figure 1A** and **C**). In addition to dendritic characteristics, we measured other anatomical features of MN24. The cell body of MN24 is located outside of the neuropile at 25.6 ± 0.8 μm from the midline (**Figure 1A**). The axonal process of MN24 extends 7.9 ± 0.8 μm towards the midline before diverging away from it (**Figure 1A**), forming a ‘routing’ pattern. Following this divergence, the axon exits the CNS and then innervates the target muscle 24 (**Figure 1A**). Additionally, we quantified the area encompassed by the axonal routing, finding it to be 36.2 ± 3.6 μm^2^ (**Fig. 2** for details of area measurement). Together, the arrangement of MN24 neuronal processes—precisely, its dendrite formation—is stereotypically positioned, suggesting MN24 dendrites are regulated in a spatial manner.

**Figure 1:**
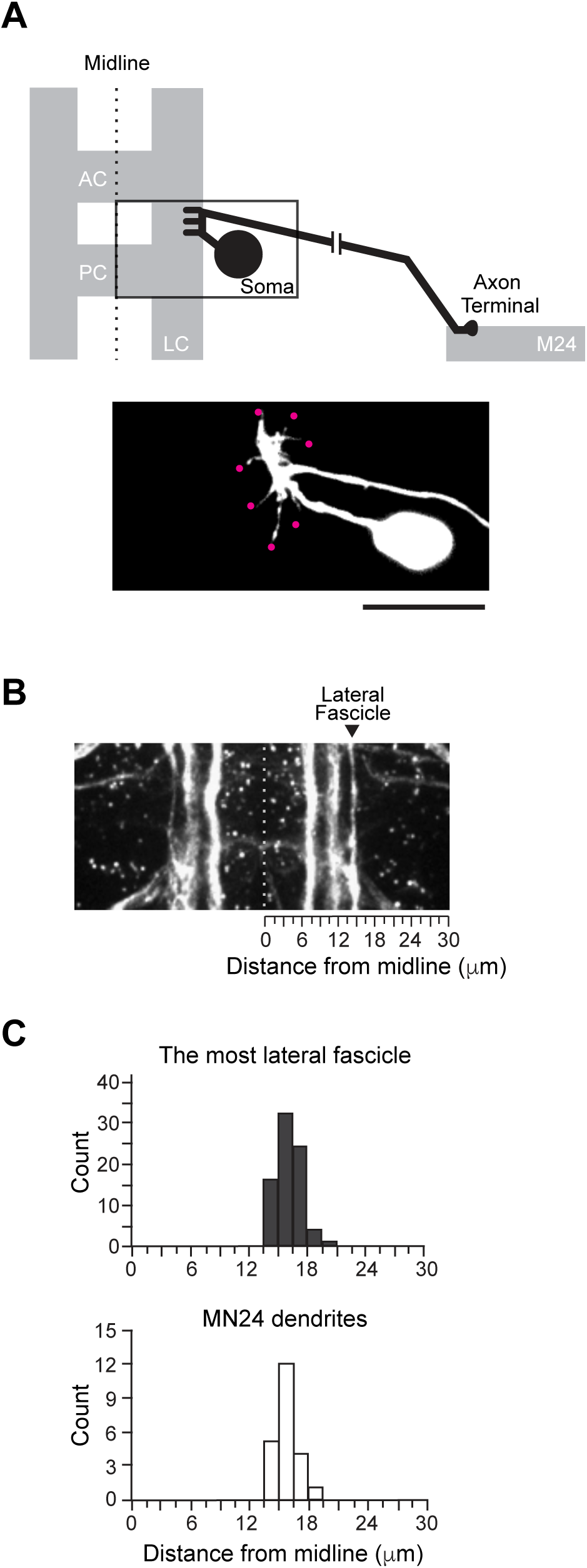
Neuronal Fascicle Spatially Aligns with the Position of MN24 Dendrite Formation. (**A**) Top panel shows a schematic of MN24 (black) within the ventral nerve cord of an embryo. The axon stereotypically projects out of the soma, anteriorly along the edge of the longitudinal connective (LC), and away from the midline to target muscle 24 (M24). The bottom panel shows a representative fluorescence image of a lipophilic-dye-labeled MN24. At 15:00h AEL, MN24 form their dendritic processes (magenta dots) at stereotyped positions on the axon routing. For all subsequent images, anterior is to the top, and medial is to the left. AC: Anterior commissure. PC: Posterior commissure. Scale bar, 10 μm. (**B**) Representative fluorescence images of FasII-positive longitudinal fascicles within the ventral nerve cord. The stereotyped most lateral FasII-positive fascicle structure (arrowhead) provides a frame of reference to characterize the mediolateral position of MN24 dendrites. Gray dashed line depicts the midline. (**C**) Distribution plots of the mediolateral positions of MN24 dendritic branches (white) (n = 22 neurons) and FasII-positive lateral fascicle (dark gray) (n = 77 hemisegments), where 0 μm indicates the distance from the CNS midline.

**Figure 2:**
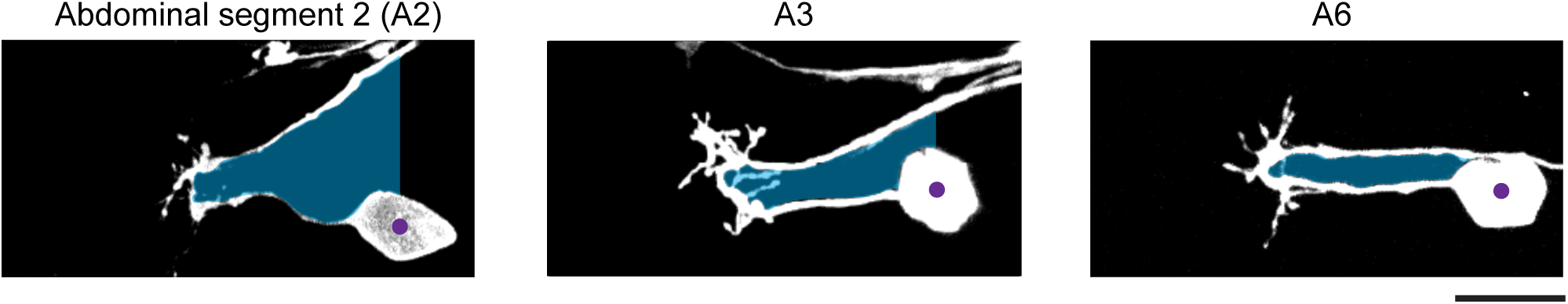
Segment-specific MN24 morphologies in the Wild-Type Background. Representative images depicting the morphology of wild-type MN24 in different abdominal segments are shown. These images show the characteristic dendrites and axon routing, observed in Figure 1A. Notably, the angle of the axon segment projecting towards the muscle varies in a segment-specific manner. Axon routing area (shaded blue) is measured as the area within the loop. For “open” axon routing areas, (left and middle panels), we define the center of the cell body (purple dot) and use the perpendicular line to the soma center as the border for measurement of the axon routing area. Scale bar, 10 μm.

### Potential Guiding Role of the Most Lateral Fascicle Structure to MN24 Dendritogenesis

The emergence of primary dendritic branches of MN24 at specific lateral positions within the CNS prompts the following question: What spatial cue guides MN24 to generate its branches at the precise location? To understand the positioning of MN24 dendrites relative to established positional landmarks, we performed immunostaining on wild-type embryos using an anti-FasII antibody. This antibody reveals a set of axon tracts, each forming distinct longitudinal fascicles within the neuropile (Landgraf et al., 2003a). These tracts run along the anterior-posterior axis and parallel to each other in the mediolateral direction (**Figure 1B**). Notably, the most lateral fascicle is located 16.2 ± 0.1 μm from the midline, closely mirroring the positioning of MN24 dendrites at 15.8 ± 0.3 μm (**Figure 1C**). Due to the incompatibility between immunohistochemistry and lipophilic dye labeling techniques, as detergent washes away the dye, we were unable to simultaneously image their structures. However, our quantitative analysis indicates their proximity, suggesting that the lateral fascicle might play a crucial positional role in MN24 dendritogenesis.

### Loss of Dscam1 Disrupts MN24 Dendritic Processes

As previously demonstrated (Kamiyama et al., 2015), the *dscam1* gene plays a prominent role in the outgrowth of primary dendritic branches in the aCC motoneuron, evidenced by the near elimination of these branches in the *dscam1* null mutants (*dscam1^-/-^*). Building upon this, we investigated whether *dscam1* similarly regulates dendritogenesis in MN24. We characterized the dendritic outgrowth of MN24 in embryos homozygous for the *dscam1* mutation. In *dscam1^-/-^*, we observed a significant decrease in the number of primary dendritic branches. On average, there were 1.3 ± 0.4 branches in *dscam1^-/-^*compared to the wild-type, which had 7.6 ± 0.3 (**Figure 3A-B**). Interestingly, in the mutant background, we observed a notable ‘collapse’ in the axonal routing of MN24 (**Figure 3A**). On average, the area of the axonal routing in the mutants was significantly reduced, measuring only 4.1 ± 4.4 μm^2^, in contrast to the wild-type area, which was 36.2 ± 3.6 μm^2^ (**Figure 3C**). Further cellular characterization in the *dscam1^-/-^*mutant revealed that while MN24 extends its axon around the target muscle region in most cases, there were occasional instances where it failed to specifically reach the target muscle 24 (**Figure 3D-E**). Despite these minor defects, the overall pattern of axon guidance in MN24 remains intact. These results suggest that the loss of *dscam1* specifically impacts the development of primary dendritic branches and the routing of the axon shaft in MN24 before exiting the VNC.

**Figure 3:**
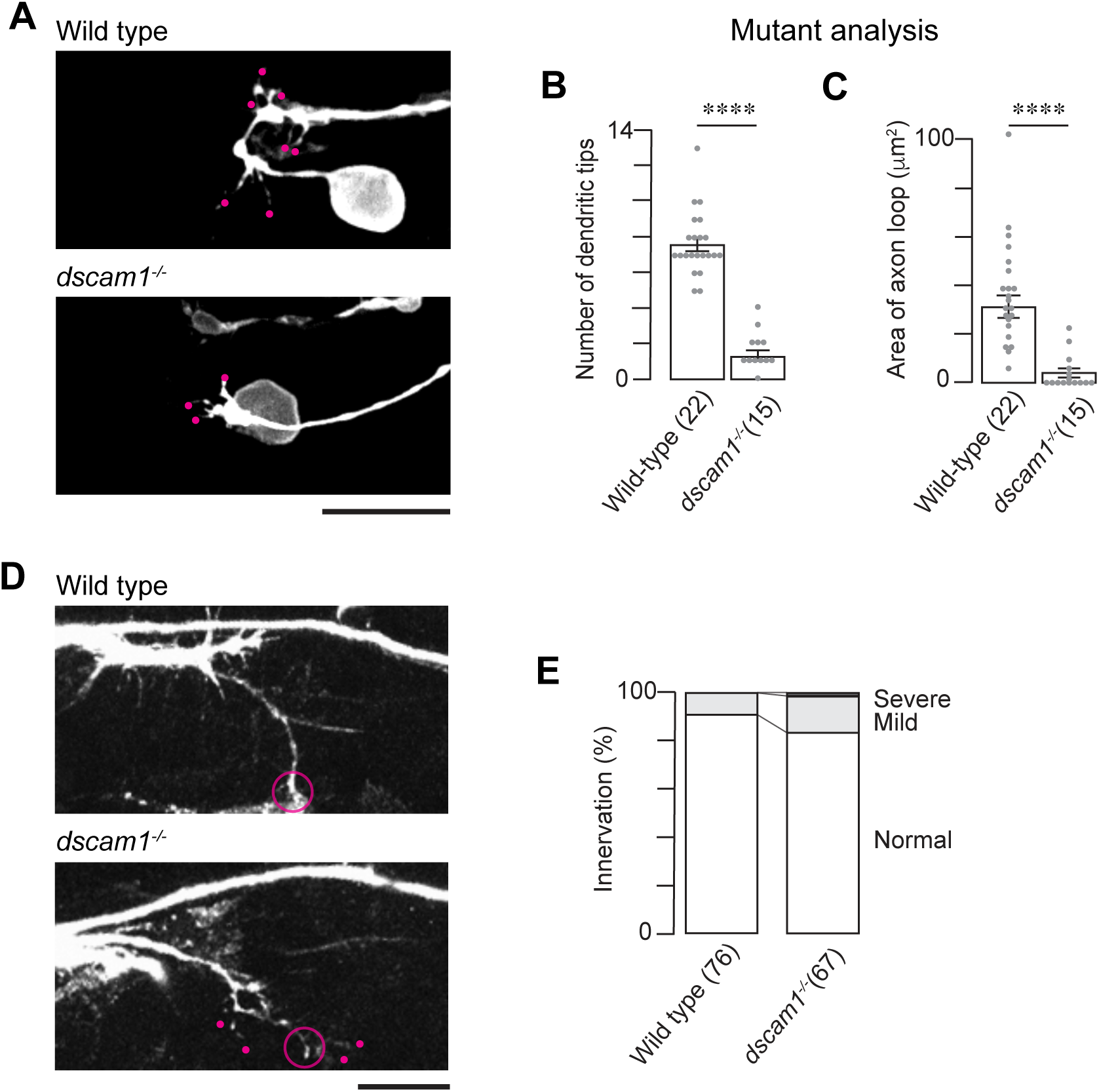
*dscam1* is Required for MN24 Neurite Development. (**A**) Representative fluorescence images of MN24 within wild-type (top panel) and *dscam1^-/-^*mutant (bottom panel) backgrounds. (**B** and **C**) Comparison of mean primary dendritic branch numbers (**B**) and axon routing areas (**C**) within wild-type and *dscam1^-/-^* mutant backgrounds; using Mann–Whitney U test. For all graphs, the sample size of neurons is denoted by the number in the parentheses of each genotype unless otherwise specified. For all subsequent statistical analyses, symbols indicate the following: ****, *p* < 0.0001; ***, *p* < 0.001; **, *p* < 0.01; *, *p* < 0.05; ns – not significant. (**D**) Immunofluorescence staining of FasII at 15:00h AEL shows the visual comparison between axon terminals in wild-type (top panel) and *dscam1^-/-^*mutant (bottom panel) backgrounds. Representative image displaying FasII staining in the wild-type and mutant backgrounds exhibits innervation by the SNa nerve branch (open circle). However, in the *dscam1^-/-^*mutant background, the SNa sub-branches have some mild targeting defects (white dots). (**E**) Quantification of SNa innervation defects within wild-type (n = 76 hemisegments) and *dscam1^-/-^* mutant (n = 67 hemisegments) backgrounds. Data is represented as a percentage – number of hemisegments with innervation defects over the total number of hemisegments observed. SNa innervation defects were characterized as mild (light gray) or severe (dark gray) when the SNa sub-branch had targeting defects or the SNa branch did not exit the nerve cord, respectively. Scale bars, 10 μm in (**A** and **D**).

Since we hypothesize that the most lateral FasII-positive fascicle might be involved in MN24 dendritogenesis, it is crucial to assess its phenotype in the mutant. Following staining of the *dscam1^-/-^* mutant with the anti-FasII antibody, we observed thinning in the lateral fascicle and, on some occasions, a ‘wavy’ pattern. However, for the most part, the mutant lateral fascicle appeared relatively normal, where 87.1% of mutant fascicles from the 66 observed hemisegments contained no breakage similar to 89.0% of those from wild-type containing no breakage. (**Figure 4A-B**). Based on these observations, we anticipate that the close proximity between this fascicle and MN24 is largely maintained.

**Figure 4:**
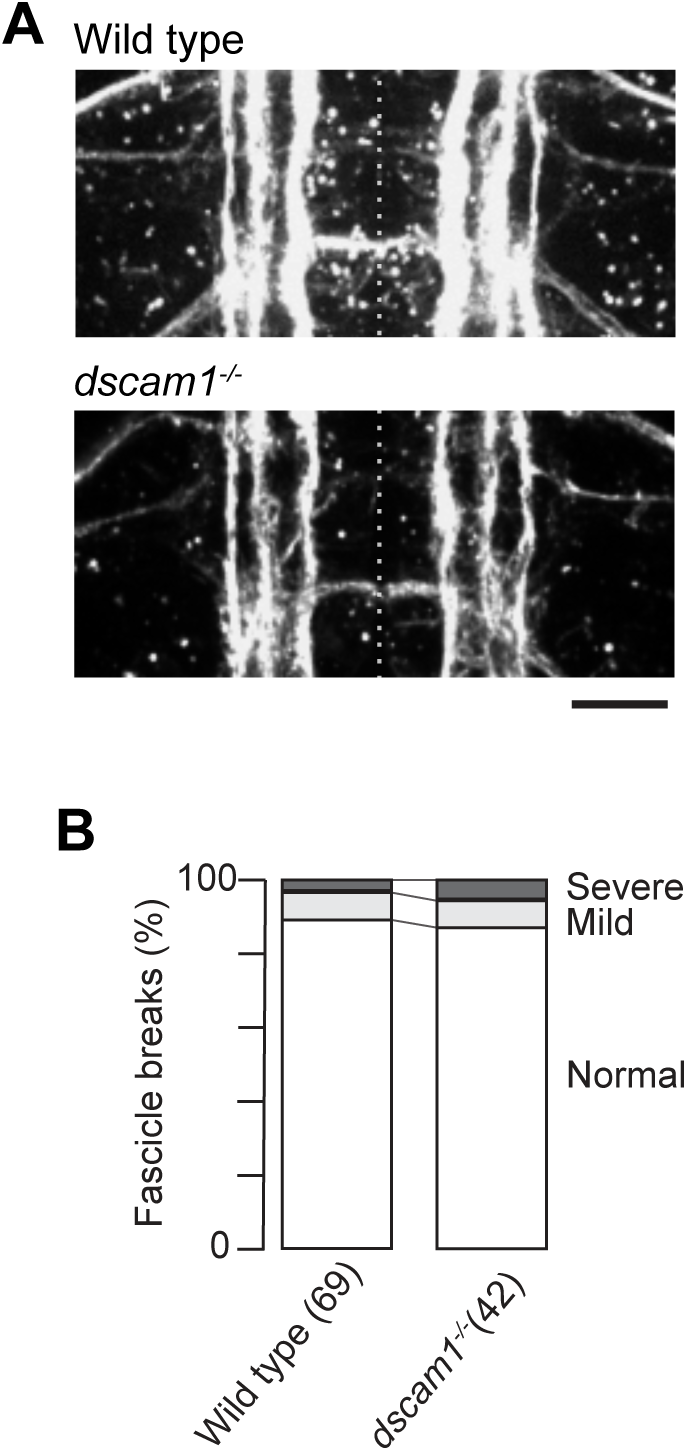
Longitudinal Fascicles in Wild-Type and *dscam1^-/-^* Mutant Backgrounds. (**A**) Representative fluorescence images of FasII-positive axon tracts within wild-type (top panel) and *dscam1^-/-^* mutant (bottom panel) backgrounds. Scale bar, 10 μm. (**B**) Quantification of lateral fascicle defects within wild-type (n = 77 hemisegments) and *dscam1^-/-^* mutant (n = 66 hemisegments) backgrounds. Data is represented as a percentage – length of lateral fascicle defects over total lateral fascicle length. Lateral fascicle defects were characterized as mild or severe when the lateral fascicle was thinning or contained a break, respectively.

In conclusion, our findings indicate that Dscam1 may act as a crucial positional cue for dendritic outgrowth and axonal routing in MN24. This aligns with our previous observations showing a high concentration of Dscam1 proteins at the neuropile, the site of MN24 dendritogenesis and axonal routing (Kamiyama et al., 2015). However, the exact mechanism—whether these defects in MN24 are a direct result of *dscam1* loss specifically in MN24 or a secondary effect arising from a global loss of *dscam1*—remains to be elucidated.

### Dual Roles of *dscam1* in Dendritic Outgrowth and Axonal Routing in MN24

To further elucidate the mechanism, we conducted cell-type specific manipulation of *dscam1* using a short hairpin RNA (shRNA) targeting *dscam1* under *UAS* control for gene knockdown (*UAS-dscam1 RNAi*). The efficacy of this *UAS-dscam1 RNAi* line, previously validated (Kamiyama et al., 2015), was apparent when expressed under the control of the pan-neuronal GAL4 driver, *elav-GAL4*, leading to the elimination of Dscam1 proteins from the embryonic CNS. Crossing this RNAi line with *hh-GAL4*, we selectively knocked down *dscam1* in MN24. This targeted approach resulted in a significant reduction in dendritic branches—on average, MN24-specific RNAi knockdown exhibited only 1.7 ± 0.4 primary dendritic branches, compared to control embryos, which had 6.8 ± 0.5 (**Figure 5A-B**). Notably, reintroducing a single isoform of *dscam1* (*UAS-dscam1^exon^ ^17.2^*) into MN24 in the *dscam1* mutant background did not restore the normal dendritic count (2.3 ± 0.5 for rescue and 1.1 ± 0.7 for mutant control) (**Figure 5C-D**).

**Figure 5:**
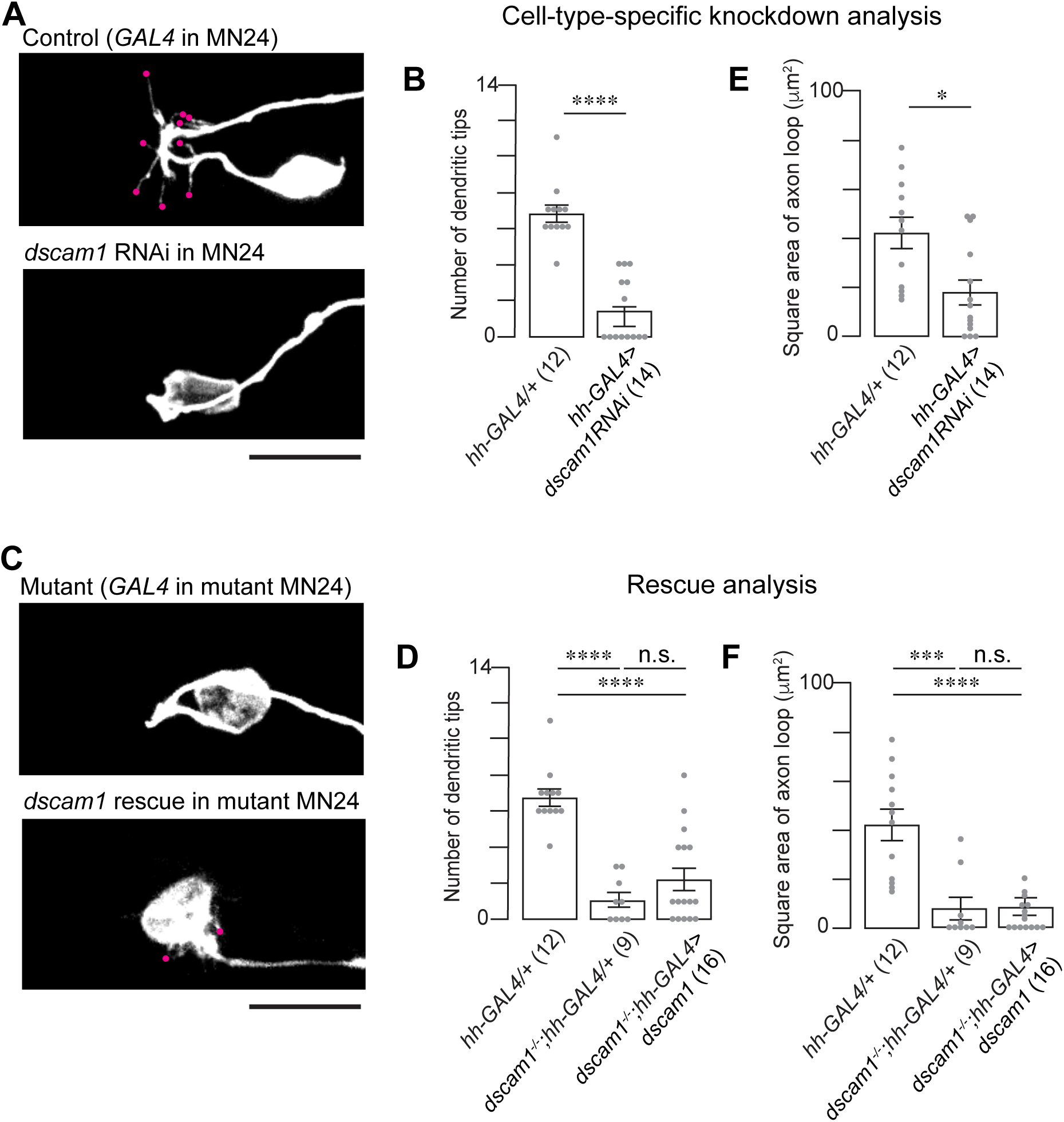
Dscam1 Plays a Cell-Autonomous Role for MN24 Neurite Development. (**A**) Representative fluorescence images of MN24 in wild-type background expressing *hh*-GAL4 driver (top panel) and *dscam1* RNAi expressed under the control of the *hh*-GAL4 driver (bottom panel). (**B** and **E**) Comparison of mean primary dendritic branch numbers (**B**) and axon routing areas (**E**) of MN24 in wild-type background expressing *hh-*GAL4 driver and *dscam1* RNAi expressed under the control of the *hh*-GAL4 driver; using Mann–Whitney U test. (**C**) Representative fluorescence images of MN24 in *dscam1^-/-^* mutant background expressing *hh*- GAL4 driver (top panel), and *dscam1^-/-^* mutant background resupplied *dscam1* expressed under the control of the *hh*-GAL4 driver (bottom panel). (**D** and **F**) Comparison of mean primary dendritic branch numbers (**D**) and axon routing areas (**F**) of MN24 in wild-type background expressing *hh*-GAL4 driver, *dscam1^-/-^* mutant background expressing *hh*-GAL4 driver, and *dscam1^-/-^* mutant background resupplied *dscam1* expressed under the control of the *hh*-GAL4 driver; using Kruskal–Wallis test followed by Dunn’s multiple comparisons test. Scale bars, 10 μm in (**A** and **C**).

Regarding axonal routing, the MN24-specific RNAi knockdown of *dscam1* partially replicated the knockout phenotype. Knocking down *dscam1* led to alterations in the axonal routing of MN24, with the average routing area measuring 14.0 ± 4.8 μm^2^, compared to the control’s 42.0 ± 5.5 μm^2^ (**Figure 5A** and **E**). However, this phenotype was less severe than in the knockout control, which had an average loop area of 7.7 ± 5.7 μm^2^ (see also **Figure 5F**, the second bar). Reintroducing the *dscam1* gene into MN24 in the *dscam1^-/-^* background did not significantly rescue the axonal routing structure observed (8.5 ± 4.2 μm^2^ for rescue) (**Figure 5B** and **F**).

From these findings, we draw two conclusions: (1) the RNAi knockdown results suggest that *dscam1* serves a cell-autonomous function in the dendritogenesis and axonal routing of MN24, and (2) the rescue results indicate that *dscam1* alone is not sufficient for the formation of both cellular structures in MN24. Furthermore, these results imply the possibility that *dscam1*, when expressed in other cells, contributes to MN24 morphogenesis, indicating a non-cell-autonomous function of dscam1 in these processes.

### Scp2-GAL4: Enabling Selective Expression of Transgenes in Lateral Fascicles

To elucidate the non-cell-autonomous functions of *dscam1*, we considered that the most lateral FasII-positive fascicle might provide positional cues to MN24, potentially mediated by Dscam1. To test this hypothesis, we must manipulate the *dscam1* gene in the lateral fascicle. However, due to the absence of a reported GAL4 line specifically labeling the most lateral fascicle, we embarked on a screening to identify a new GAL4 driver. By crossing approximately 20 GAL4 lines with *UAS-mCD4-tdGFP*, we identified a promising candidate, *Scp2-GAL4*. This GAL4 line labels a subset of interneurons that contribute to the formation of the most lateral fascicle. The expression pattern observed in *Scp2-GAL4* highlights neuronal processes from interneurons, segregated into either the medial or lateral fractions of the FasII-positive fascicles (**Figure 6**). Additionally, this GAL4 line targeted aCC and RP2 motoneurons in 40.6% and 34.3% of the observed hemisegments (n=32), respectively. MN3 and MN19 were also labeled, though less frequently, at 6.3% for each of the hemisegments observed. Importantly, *Scp2-GAL4* does not label MN24 or any related motoneurons within the same SNa nerve tract. In conclusion, we identified *Scp2-GAL4* as a GAL4 line that facilitates the expression of *UAS* transgenes in two of the FasII-positive fascicles, notably including the most lateral fascicle.

**Figure 6:**
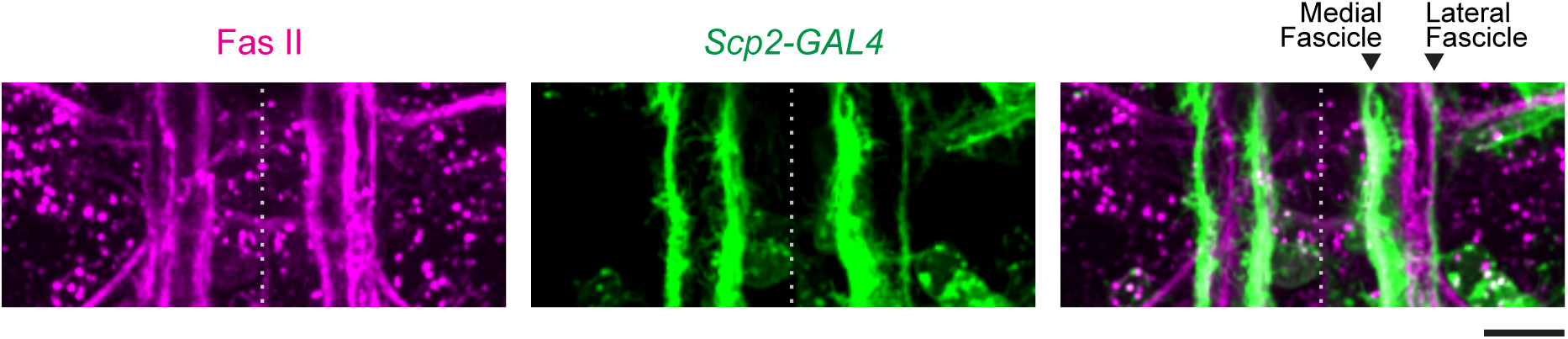
*Scp2*-GAL4 Driver Allows Labeling of Lateral Fascicle. Representative images of neuronal fascicles labeled by membrane-bound GFP under the control of *Scp2*-GAL4 driver (green) or immunostained with anti-FasII antibody (magenta). *Scp2*-positive fascicles include the medial and lateral fascicles (arrowheads) and exclude the intermediate fascicle. Scale bar, 10 μm.

### Dscam1 in the Lateral Fascicle is Necessary for MN24 Dendritogenesis and Axonal Routing

Using the *Scp2-GAL4* driver, we simultaneously implemented *UAS-dscam1 RNAi* for specific gene knockdown and *UAS-mCD4-tdGFP* for targeted cell labeling. This approach led to a significant reduction in dendritic branches—on average, MN24 exhibited 2.0 ± 0.4 primary dendritic branches, compared to the control, which had 8.3 ± 0.6 (**Figure 7A-B**). These results strongly support the concept of a non-cell-autonomous function for *dscam1*. Interestingly, subsequent attempts to rescue the dendritic phenotype by reintroducing *dscam1* into Scp2-positive neurons were unsuccessful in reversing the mutant phenotype in MN24 (1.4 ± 0.5 for rescue and 2.1 ± 0.5 for mutant control) (**Figure 7C-D**). This suggests that the expression of the *dscam1* gene only in Scp2-positive neurons is not sufficient for MN24 dendritogenesis.

**Figure 7:**
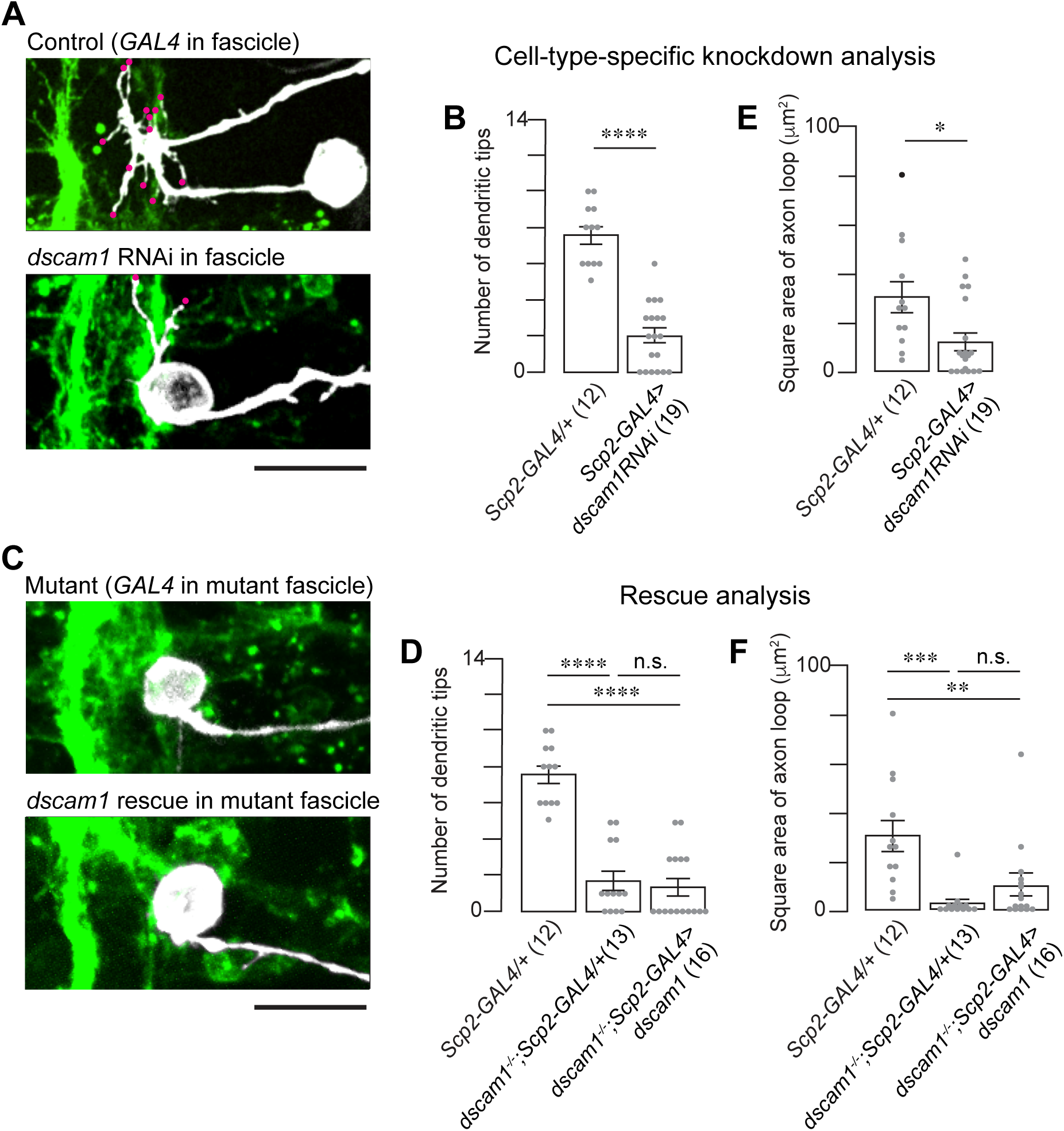
*Scp2*-Positive Lateral Fascicle Provides Non-Cell-Autonomous Dscam1 for MN24 Neurite Development. (**A**) Representative fluorescence images of MN24 in wild-type background expressing *Scp2*-GAL4 driver (top panel) and *dscam1* RNAi expressed under the control of the *Scp2*-GAL4 driver (bottom panel). (**B** and **E**) Comparison of mean primary dendritic branch numbers (**B**) and axon routing areas (**E**) of MN24 in wild-type background expressing *Scp2*-GAL4 driver and *dscam1* RNAi expressed under the control of the *Scp2*-GAL4 driver; using Mann–Whitney U test. (**C**) Representative fluorescence images of MN24 in *dscam1^-/-^* mutant background expressing *Scp2*-GAL4 driver (top panel), and *dscam1^-/-^* mutant background resupplied *dscam1* expressed under the control of the *Scp2*-GAL4 driver (bottom panel). (**D** and **F**) Comparison of mean primary dendritic branch numbers (**D**) and axon routing areas (**F**) of MN24 in wild-type background expressing *Scp2*-GAL4 driver, *dscam1^-/-^*mutant background expressing *Scp2*-GAL4 driver, and *dscam1^-/-^*mutant background resupplied *dscam1* expressed under the control of the *Scp2*-GAL4 driver; using Kruskal–Wallis test followed by Dunn’s multiple comparisons test. Scale bars, 10 μm in (**A** and **C**).

Similarly, RNAi knockdown of *dscam1* using *scp2-GAL4* led to a ‘collapse’ in the axonal routing of MN24. The average axon routing area was measured at 12.6 ± 4.6 μm^2^, significantly reduced compared to the control, which was measured at 35.6 ± 5.8 μm^2^ (**Figure 7A** and **E**). Additionally, when we resupplied the *dscam1* gene only to Scp2-positive neurons in *dscam1^-/-^*, there was no observed rescue of the axonal routing structure (9.4 ± 4.1 μm^2^ for rescue and 2.4 ± 4.0 μm^2^ for mutant control) (**Figure 7B** and **F**). Importantly, upon imaging in the rescue experiments, we found that the cell bodies of MN24 were variably positioned relative to the most lateral fascicles; mutant MN24 had a cell body position that seemed more medially shifted compared to control MN24 (for example, see **Figure 7A and 8A**). Inspired by this observation, we measured the positions of both the cell bodies and the fascicle relative to the midline. We discovered that while the position of the most lateral fascicle remained unchanged (15.3 ± 0.4 μm for mutant and 14.3 ± 0.5 μm for control), the cell bodies of MN24 were differently positioned, often closer to the lateral fascicle (19.8 ± 0.9 μm for mutant and 26.7 ± 1.0 μm for control) (**Figure 8B-C**). This led us to speculate that the reduced area of the axon loop might be a secondary defect—due to the proximity of MN24 cell bodies to the most lateral fascicle, there may be insufficient space for the axonal routing to form properly in these genetic backgrounds (see the Discussion).

**Figure 8:**
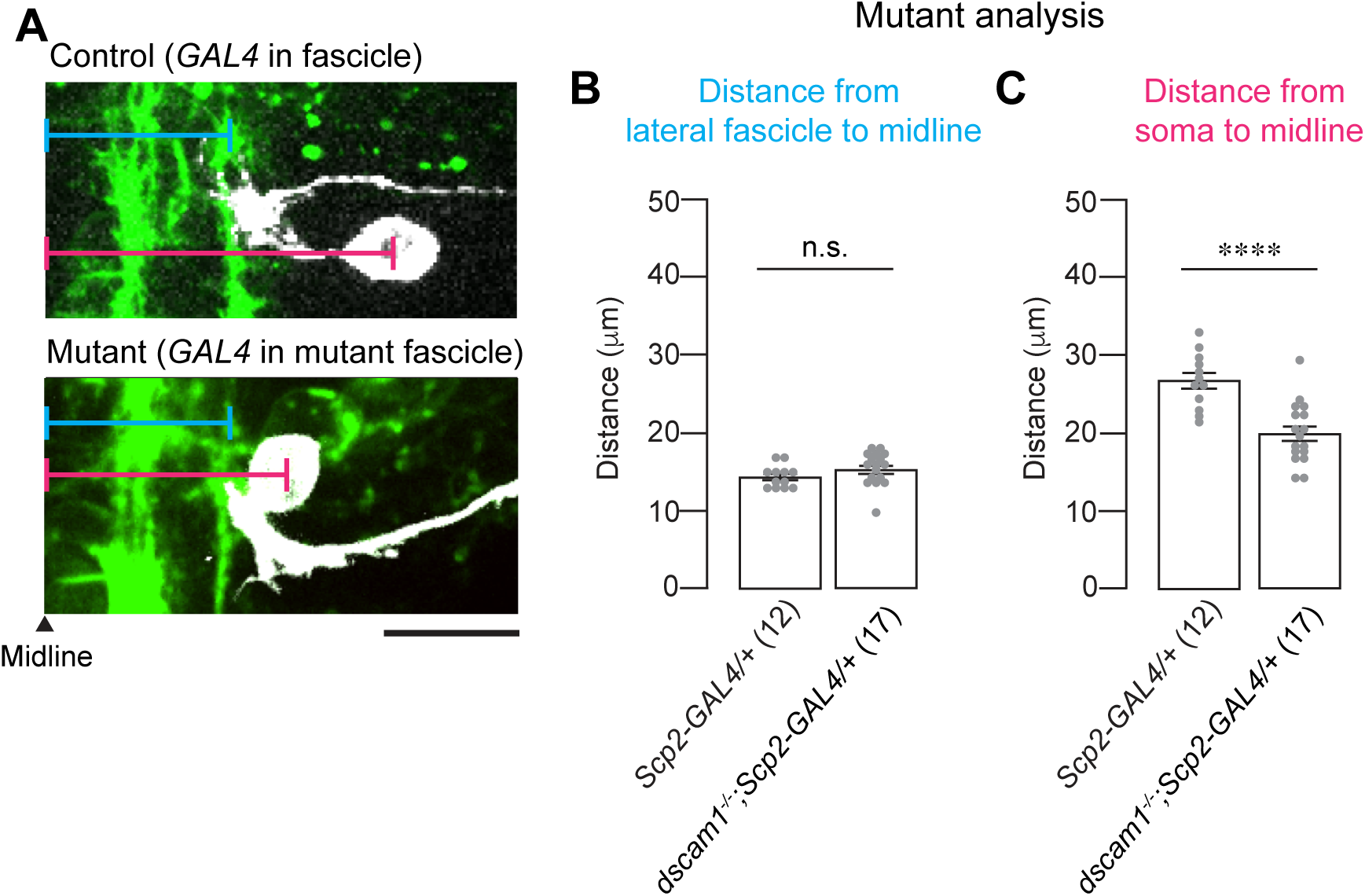
MN24 Soma Position is Medially Shifted in the *dscam1^-/-^* Mutant Background. (**A**) Representative images of MN24 at 15:00h AEL in wild-type background expressing *Scp2*-GAL4 driver (green) (top panel) and *dscam1^-/-^* mutant background expressing *Scp2*-GAL4 driver (bottom panel). Blue and pink bars indicate the distance (μm) from the lateral fascicle and soma, respectively, to the midline. Scale bar, 10 µm. (**B**) Quantification of lateral fascicle position in wild-type background expressing *Scp2*-GAL4 driver and *dscam1^-/-^* mutant background expressing *Scp2*-GAL4 driver; using Welch’s *t* test. The *Scp2*-positive lateral fascicle does not have a mediolateral shift in the *dscam1^-/-^* mutant background. (**C**) Quantification of MN24 soma position in wild-type background expressing *Scp2*-GAL4 driver and *dscam1^-/-^* mutant background expressing *Scp2*-GAL4 driver; using Welch’s *t* test. MN24 soma in the *dscam1^-/-^* mutant background expressing *Scp2*-GAL4 driver has a more medial shift compared to that of the wild-type background.

### Dscam1 Mediates Interaction Between MN24 and the Lateral Fascicle for Proper Dendritogenesis and Axonal Routing of MN24

Our experiments indicate that both cell-autonomous and non-cell-autonomous functions of *dscam1* are essential for dendritogenesis and axonal routing in MN24 and suggest that Dscam1 on either side of the neuronal membranes function to guide MN24 neurite processes. If Dscam1 serves as a positional cue, then we reasoned that providing Dscam1 to both MN24 and Scp2-positive neurons would restore the MN24 mutant phenotype. To directly test this hypothesis, we reintroduced *UAS-dscam1* into *dscam1^-/-^* mutants using two GAL4 drivers, *Scp2-* and *hh-GAL4*, targeting both MN24 and Scp2-positive fascicle. In alignment with our hypothesis, this dual reintroduction of *dscam1* led to a complete recovery of the MN24 dendrite count (7.4 ± 0.5 for rescue and 8.4 ± 0.6 for control) and restoration of the axonal routing structure (41.1 ± 6.2 μm^2^ for rescue and 38.6 ± 6.7 μm^2^ for control) (**Figure 9A-C**). Notably, the axonal structure recovered, with the cell bodies repositioning to locations similar to those in the controls (27.1 ± 0.9 μm for rescue and 28.6 ± 1.0 μm for control) (**Figure 10**). These findings suggest that Dscam1’s function in both MN24 and Scp2-positive neurons is crucial for dendritogenesis and axonal routing in MN24 and are consistent within the model that Dscam1 acts as a positional cue to guide MN24 development (**Figure 11A**).

**Figure 9:**
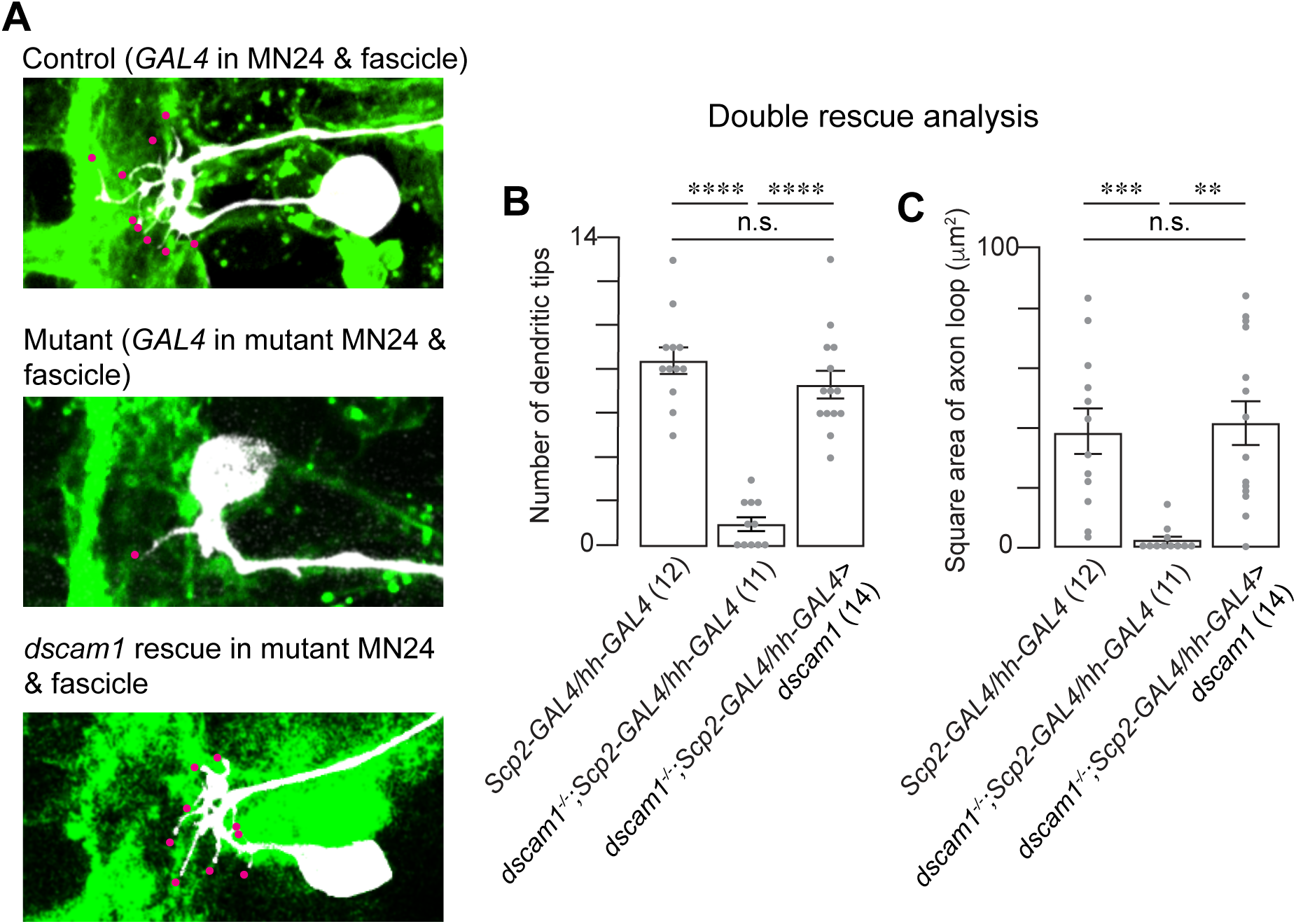
Dscam1 in Both Scp2-Positive Lateral Fascicle and MN24 is Sufficient to Restore MN24 Dendritogenesis and Axon Routing. (**A**) Representative fluorescence images of MN24 within wild-type background (top panel), *dscam1^-/-^* mutant background with combined *Scp2*- and *MN24*-specific expression of membrane-bound GFP (middle panel), and *dscam1^-/-^* mutant background with combined *Scp2*- and *MN24*-specific resupply of *dscam1* (bottom panel). Scale bar, 10 μm. (**B** and **C**) Comparison of mean primary dendritic branch numbers (**B**) and axon routing area (**C**) among MN24 in wild-type background, *dscam1^-/-^* mutant background with *Scp2*- and *MN24*-specific expression of GFP membrane-bound, and *dscam1^-/-^* mutant background with combined *Scp2*- and *MN24*-specific resupply of *dscam1*; using Kruskal–Wallis test followed by Dunn’s multiple comparisons test.

**Figure 10:**
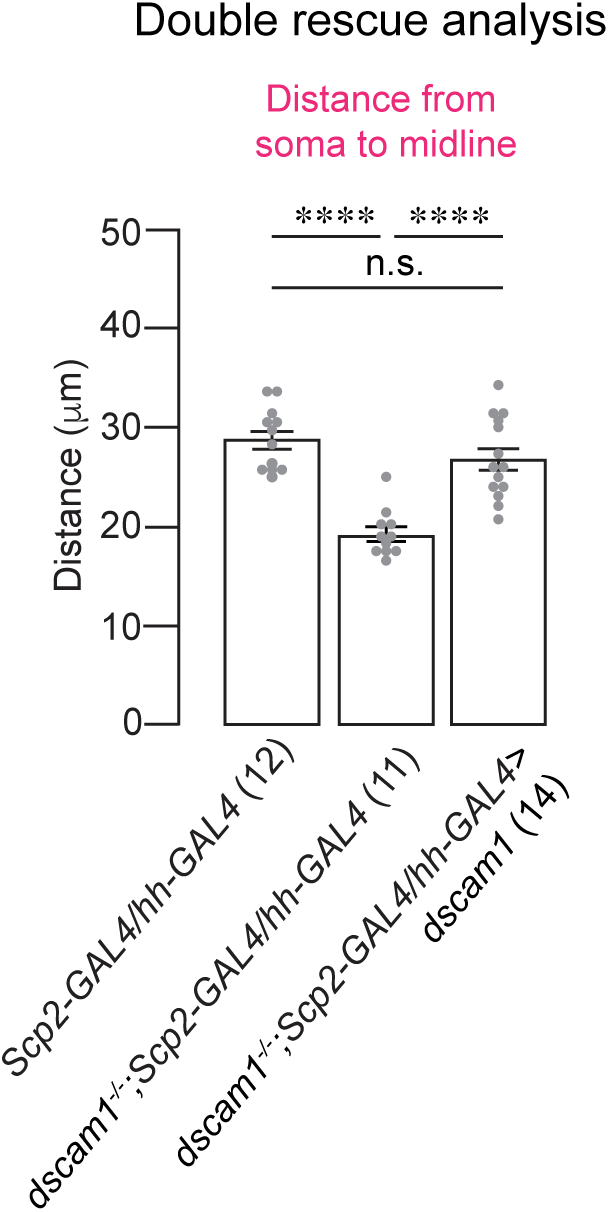
Resupplying *dscam1* in Scp2-Positive Lateral Fascicle and MN24 Restores Mutant MN24 Soma Position. Quantification of MN24 soma positions in wild-type background with *Scp2*- and *hh*-specific expression of membrane-bound GFP, *dscam1^-/-^*mutant background with *Scp2*- and *hh*-specific expression of membrane-bound GFP, and *dscam1^-/-^* mutant background with combined *Scp2*- and *hh*-specific resupply of *dscam1*; using Kruskal–Wallis test followed by Dunn’s multiple comparisons test.

**Figure 11:**
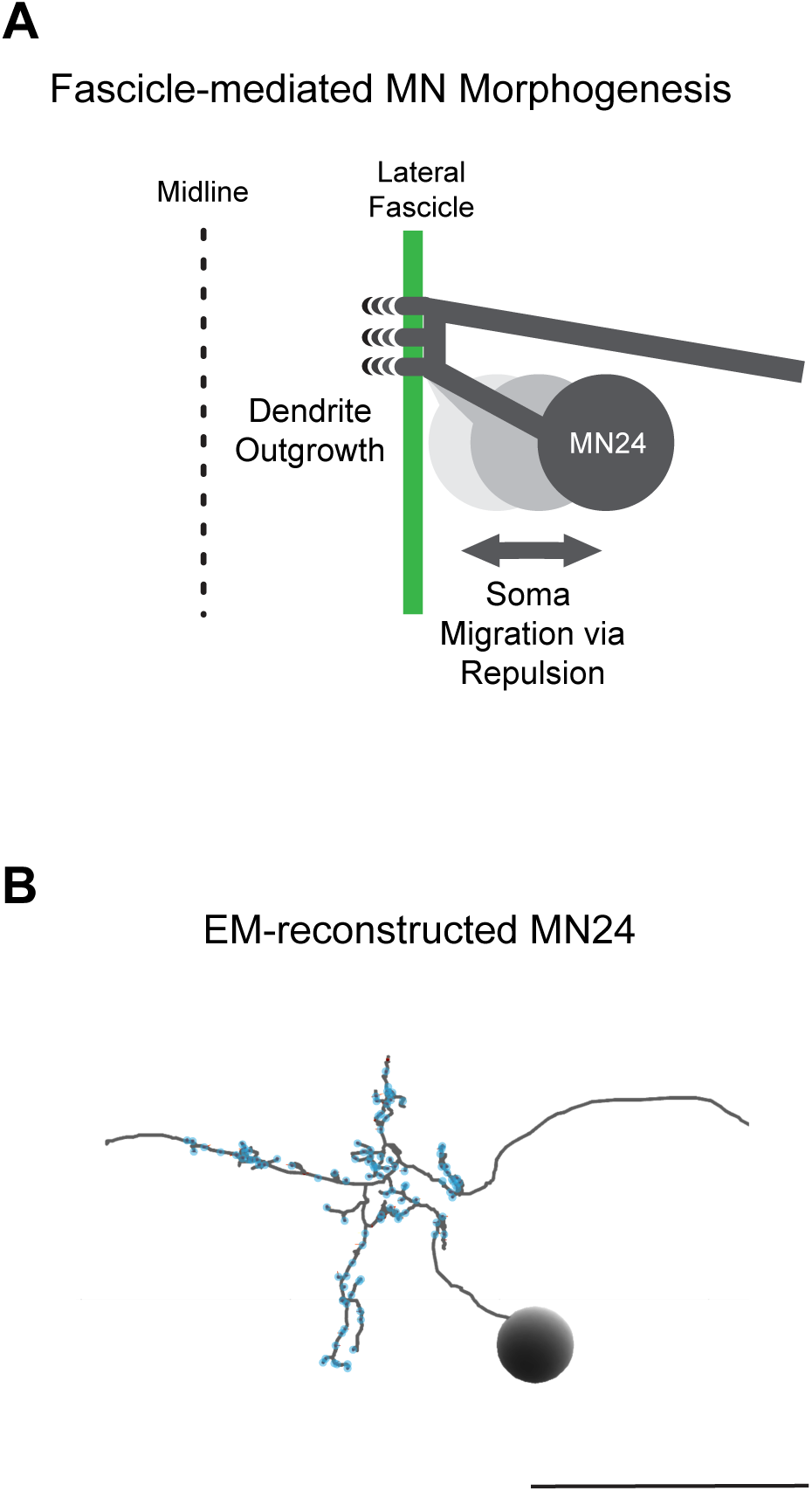
Proposed Model for Fascicle-Mediated MN24 Morphogenesis. (**A**) Schematic illustrating the proposed model of how the lateral fascicle structure mediates MN24 dendrite outgrowth and soma migration. (**B**) Electron microscopy (EM) reconstruction of a single MN23/24 in 1st instar larva. Prominent morphological structures such as dendritic outgrowth and axon routing are retained in larval MN24. The backbone is indicated by gray. Blue dots indicate synaptic sites. Scale bar, 20 μm.

## Discussion

### Dscam1 as a Positional Cue Defines the MN24 Dendritogenesis Site

Expanding on our previous findings about dendritogenesis in the aCC motoneuron (Kamiyama et al., 2015), our current study explores the function of *dscam1* during MN24 dendritogenesis. Our research presents several lines of evidence suggesting that the interaction between MN24 and Scp2-positive neurons is critical for the outgrowth of primary dendritic branches in MN24. Firstly, we demonstrate that MN24 and the Scp2-positive fascicle are in proximity. Secondly, upon knocking down *dscam1* in either MN24 or Scp2-positive fascicle, we observed a reduction in the number of primary dendritic branches, mirroring the phenotype seen in *dscam1^-/-^* mutants. This suggests that *dscam1* in both MN24 and Scp2-positive fascicle is necessary for MN24 dendritogenesis. Thirdly, our rescue experiments in *dscam1^-/-^* mutants, involving the reintroduction of *dscam1* into either MN24 or Scp2-positive fascicle, were unsuccessful. This indicates that *dscam1* function in either neuron type alone is insufficient. However, when *dscam1* was reintroduced into both MN24 and Scp2-positive fascicle in *dscam1^-/-^* mutants, we could fully rescue the dendritic phenotype. Finally, the observation that MN24 axons still contact the Scp2-positive fascicle in *dscam1^-/-^*mutants rules out the possibility that the reduced number of dendrites is due to a mislocation of these neural processes. Consequently, we propose that Dscam1 provides a positional cue for MN24 through cell-cell contact, defining the site of dendritic outgrowth (**Figure 11A**). This mechanism echoes how vertebrate DSCAM guides retinal ganglion cell (RGC) dendrites and bipolar cell axons for synapse formation in the chick retina (Yamagata and Sanes, 2008), suggesting a potentially conserved principle in dendritic outgrowth mediated by inter-neuronal Dscam1 interactions.

Our study specifically targets the early developmental stage of the MN24, around 15:00 after egg laying (AEL), a pivotal time when primary dendritic branches start to emerge. The critical role of these initial branches in forming the foundation for higher-order branches and synaptic formations has yet to be fully established, but emerging data offer promising insights. Recent advancements in comprehensive connectome efforts have facilitated the reconstruction of the entire CNS in the first instar larva, and this valuable data is accessible in a publicly available database (Saalfeld et al., 2009; Winding et al., 2023). Using this resource, we have examined an electron microscopy (EM) reconstructed model of MN24. This model reveal that the higher-order branches are situated in the regions of the axonal turning points, corresponding to the area where MN24’s primary dendrites initiate (**Figure 11B**). Notably, these branches exhibit numerous synapses throughout their structure. This observation suggests a developmental progression from primary dendritic branches to the establishment of functional synapses in MN24.

### The Roles of Dscam1 in Axonal Routing and Soma Migration of MN24

In addition to the observed loss-of-dendrite phenotype, our study has revealed axonal routing defects in MN24 in *dscam1^-/-^* mutants. Normally, MN24 axons project ventrally, reaching the most lateral FasII-positive fascicle, and then undergo a crucial lateral turn as part of their axonal routing process. However, in *dscam1^-/-^* mutants, this axonal routing is notably compromised. One potential explanation for this diminished axonal routing in *dscam1^-/-^*mutants could be related to a migration defect of the soma in MN24. In *dscam1^-/-^* mutants, the soma position is observed to be closer to the lateral fascicle (**Figure 7C** and **Figure 8A** and **C**). Interestingly, reintroducing *dscam1* into both MN24 and Scp2-positive neurons corrects the soma’s position, subsequently leading to the restoration of the normal axonal loop structure (**Figure 9C** and **Figure 10**).

The role of *dscam1* in soma migration during brain development in *Drosophila* is an emerging research interest. A critical study by Liu et al. focusing on larval medulla neurons has provided significant insights into this process (Liu et al., 2020). They revealed that within the fly visual system, the cell bodies of sister neurons from the same lineage exhibit mutual repulsion. This event contributes to the formation of columnar structures. This repulsive interaction is mediated by inter-neuronal interaction between Dscam1. Inspired by these findings, we propose a similar mechanism in the MN24 system. We propose a model where Dscam1 orchestrates a repulsive interaction between the soma of MN24 and the Scp2-positive fascicle (**Figure 11A**). Given the lateral expansion of the VNC during development (Tomer et al., 2012), the soma of the MN24 is initially positioned close to the lateral fascicle and is likely to migrate laterally as development progresses. This migration would be started off by inter-neuronal Dscam1 interactions.

### Single Isoform of Dscam1 for Rescue in Morphological Defects in MN24

An impressive diversity of 19,008 isoforms, each with different extracellular domains, can arise from the *dscam1* gene through alternative splicing of three variable exon clusters (Schmucker et al., 2000; Tomer et al., 2012; Sun et al., 2013). These extracellular domains can bind in a homophilic and isoform-specific manner (Wojtowicz et al., 2004; Wojtowicz et al., 2007). Intriguingly, each neuron in the fly is found to express a distinct and limited set of Dscam1 isoforms (Hattori et al., 2007; Hattori et al., 2009). Consequently, the isoform-specific binding characteristics of Dscam1 facilitate homophilic repulsion exclusively among identical (or ‘self’) cells, raising questions about Dscam1 interactions between different neuron types like MN24 and Scp2-positive neurons.

In our experiments, we introduced a single isoform of *dscam1* simultaneously into different neuron types, which successfully rescued the phenotypes associated with dendritogenesis and axonal routing in MN24 (**Figure 9A-C**). These findings suggest that just one isoform of *dscam1* is sufficient for these developmental processes. This leads us to question the nature of Dscam1 trans- and homophilic interactions between different neuronal types. Several hypotheses arise: one possibility is that MN24 and Scp2-positive neurons express the same set of isoforms, potentially due to originating from the same neuronal progenitor cells, thus sharing isoform profiles. Alternatively, the trans-interaction of Dscam1 might be mediated by other molecules, forming a protein complex. For instance, in *C. elegans*, the dendritic branching of PVD neurons involves the interaction of SAX7/NMR-1 transmembrane proteins with DMA-1, mediated by the secreted LECT-2 adapter (Zou et al., 2016; Sundararajan et al., 2019). A similar mechanism might be at play in *Drosophila*, with secreted molecules (such as Slit (Dascenco et al., 2015; Alavi et al., 2016), Netrin (Andrews et al., 2008; Liu et al., 2009), or other ligands yet to be determined) bridging opposing Dscam1 membranes through their non-variable regions.

### Cross-Species Insights into DSCAM-Mediated Motor Circuit Formation

Unraveling the specific mechanisms of Dscam1 interactions among diverse neuronal types will significantly broaden our understanding of how our model generalizes to motoneurons in *Drosophila*. Additionally, the structural and functional similarities between the *Drosophila* embryonic CNS and the mammalian spinal cord highlight the potential for cross-species studies on DSCAM. The spinal cord, within the neural tube, serves as a model for axon guidance research, showcasing shared molecular mechanisms between mammals and *Drosophila* (Evans and Bashaw, 2010; Evans, 2016; Howard et al., 2019). For instance, the interaction between Netrin1 and DCC, which directs commissural axons towards the midline in mice, reflects analogous processes in the *Drosophila* embryonic CNS (Harris et al., 1996; Kolodziej et al., 1996; Mitchell et al., 1996; Fazeli et al., 1997). Recent findings from Klar’s group have significantly emphasized the role of homophilic DSCAM interactions in the fasciculation of chick commissural axons (Cohen et al., 2017). Their *in-situ* hybridization data reveal that *DSCAM* is expressed in subsets of motoneurons. Considering the close proximity of motoneuron cell bodies and dendrites to these commissural axons (Avraham et al., 2009; Avraham et al., 2010), it is plausible that axonal fascicles could influence motoneuron morphogenesis through DSCAM-mediated interactions. Future research along these lines is essential.

## Materials and Methods

### Fly Stocks

Canton-S was used as a wild-type strain (source: W. Kim). For mutant analyses, *dscam1^21^*(source: J. Wang) was used. The following lines were obtained from the Bloomington *Drosophila* Stock Center: *UAS*-*mCD4::tdGFP* (#35836), *UAS*-*mCD4::tdTomato* (#35841), *UAS*- *Dscam1 RNAi* (#38945), *hh*-GAL4 (#49437), *scp2*-GAL4 (#49538), and *Kr*-*GFP* balancer (#5195). Homozygous mutants were identified using GFP balancers. *hh-GAL4* and *Scp2-GAL4* were used for transgenic expression in MN24 and a subset of the lateral fascicle, respectively, from the Janelia GAL4 stocks. For the rescue experiments in *dscam1^-/-^*, in **Figures 3C**, **4D**, and **5A**, a single isoform of *dscam1* (*UAS-Dscam1^exon^ ^17.2^-GFP*) (source: T. Lee) was used. Specific fly genotypes in experiments are described in **Table 1**. Flies were reared at 25°C using standard procedures.

**Table 1:**
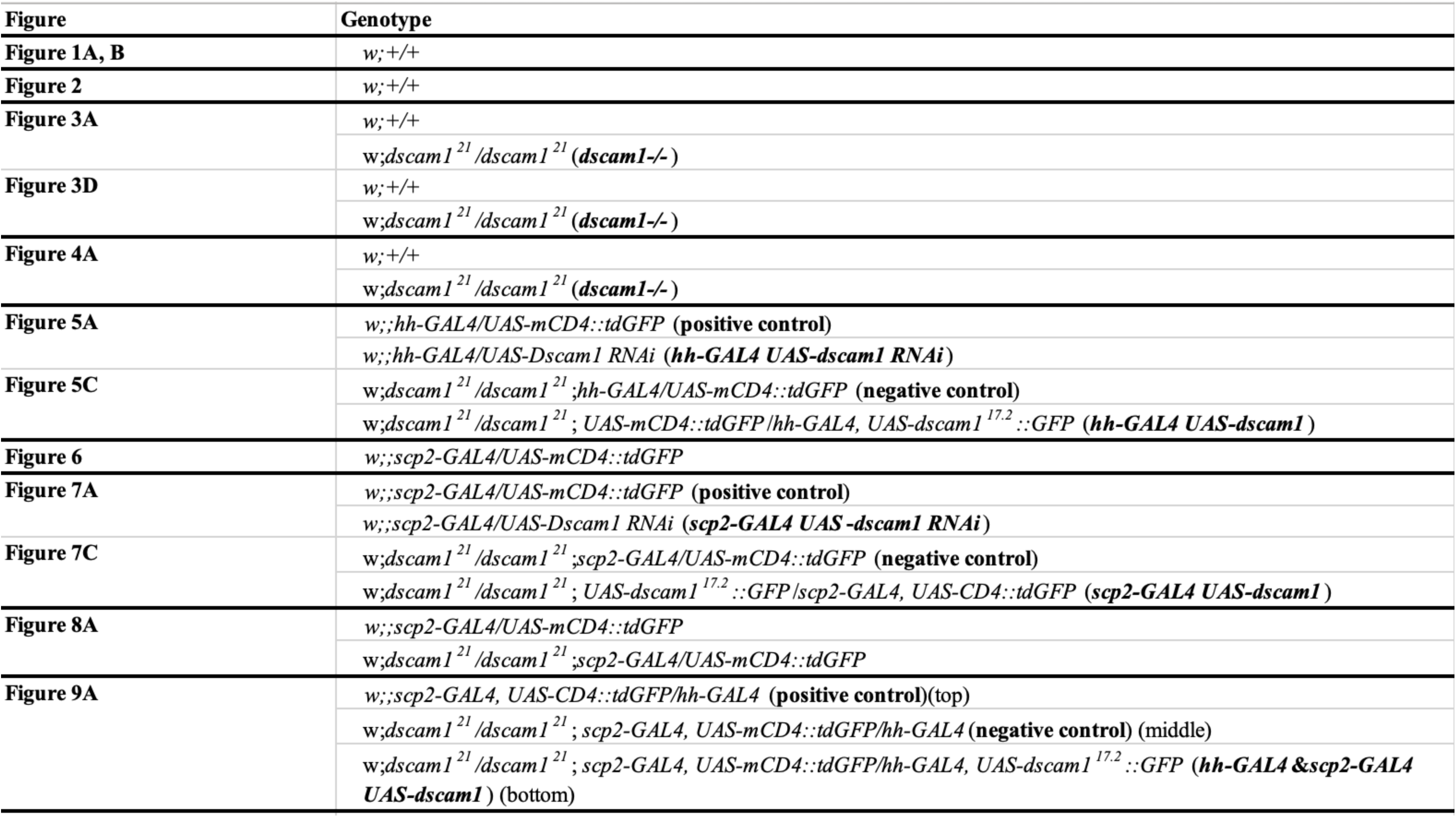
Genotypes of flies shown in this study, related to Figure 1-9.

### RNAi Experiments

For cell-specific RNAi experiments, the *UAS-shRNA* line that targets all splice variants known for *dscam1* was obtained from TRiP at Harvard Medical School via BDSC. For all examinations of *dscam1* functions, in **Figures 3A** and **4B**, the *dscam1* RNAi construct was expressed in various small subsets of neurons (*UAS-mCD4::tdTomato*/+;*UAS-dscam1RNAi*/*GAL4*). Detailed information on used *GAL4* drivers is listed in **Table 1**.

### Immunohistochemistry

Embryos were fillet-dissected, fixed with 4% paraformaldehyde for 5 min, blocked in a solution of PBS/0.01% Triton X-100 with 0.06% BSA (TBSB) for 1 hour at room temperature (RT). For labeling of Scp2-positive lateral fascicle with a reference pattern (anti-Fasciclin II [FasII] and/or anti-Horseradish Peroxidase [HRP]), the embryos were incubated with anti-FasII (mouse mAb, Developmental Studies Hybridoma Bank; 1:500) in TBSB at 4°C overnight. Samples were washed 3 x 5 min with TBSB and incubated with conjugated anti-HRP (goat mAb, JacksonImmuno; 1:500) and secondary antibodies (Alexa Fluor 647 Donkey anti-mouse, Invitrogen at 1:500) for 2 hours at RT and washed with PBS. Anti-HRP conjugation with fluorescent dyes was performed by following the same procedure as described in previous literature (Inal et al., 2021). Following immunohistochemistry, they were post-fixed with 4% paraformaldehyde for 10 min and mounted in PBS.

### Fluorescence Imaging

Confocal microscopy images of fillet embryos expressing green or red fluorescent proteins alongside far-red DiD-labeled neurons were captured using an inverted fluorescence microscope (Ti-E, Nikon) with either 40x 0.80 NA water immersion objective or 100× 1.45 NA oil immersion objective (Nikon). The microscope was attached to the Dragonfly Spinning disk confocal unit (CR-DFLY-501, Andor). Three excitation lasers (40 mW 488 nm, 50 mW 561 nm, and 110 mW 642 nm lasers) were coupled to a multimode fiber passing through the Andor Borealis unit. A dichroic mirror (Dragonfly laser dichroic for 405-488-561-640) and three bandpass filters (525/50 nm, 600/50 nm, and 725/40 nm bandpass emission wheel filters) were placed in the imaging path. Images were recorded with an electron-multiplying charge-coupled device camera (iXon, Andor).

### Labelling Dendrites and Quantifying Dendritic Processes in MN24

For phenotypic analyses of dendritic processes in wild-type and mutant backgrounds, DiD labeling (ThermoFisher) of MN24 was performed by following the same procedure as described in the literature (Inal et al., 2020). To minimize the variation in the dendritic processes in MN24 in different segments, neurons from abdominal segments 2 to 7 were imaged. Primary dendritic processes in individual MN24 that were longer than 1.0 μm were counted.

### Quantitative measurement of MN24 and Scp2-positive lateral fascicle position

MN24 was labeled with DiD and genetically encoded membrane markers (mCD4::tdGFP or mCD4::tdTomato). Similarly, Scp2-positive lateral fascicles are marked using the aforementioned genetically encoded membrane markers. Confocal stacks were acquired varying between 0.1 and 0.5 µm z-steps. The distance of the FasII- or Scp2-positive lateral fascicle from the midline was measured by first generating the FasII- or Scp2-positive lateral fascicle intensity profile perpendicular to the midline. Then, measurements are fitted in a histogram plot. Images were analyzed using Fiji (NIH). Figures were prepared using Adobe Illustrator and Photoshop.

### Experimental design and statistical analyses

A between-subject design was employed in all experiments. Immunohistochemistry and dye-labeling experiments were repeated at least two and ten times, respectively, using flies from independent crosses. Statistical analyses were performed and visualized using JMP Pro 16. The results of the statistical tests are shown in **Table 2**. All datasets were assessed for normality using Shapiro-Wilk’s test, and nonparametric tests were employed when the normality assumption was not met. Comparisons between two groups were analyzed using nonparametric Mann-Whitney U test or parametric Welch’s *t* test. Comparisons between multiples groups were analyzed using nonparametric Kruskal-Wallis test. *Post hoc* comparisons were performed using Dunn’s multiple comparison test. Error bars are shown as the Standard Error of the Mean (SEM) in the figures.

**Table 2:**
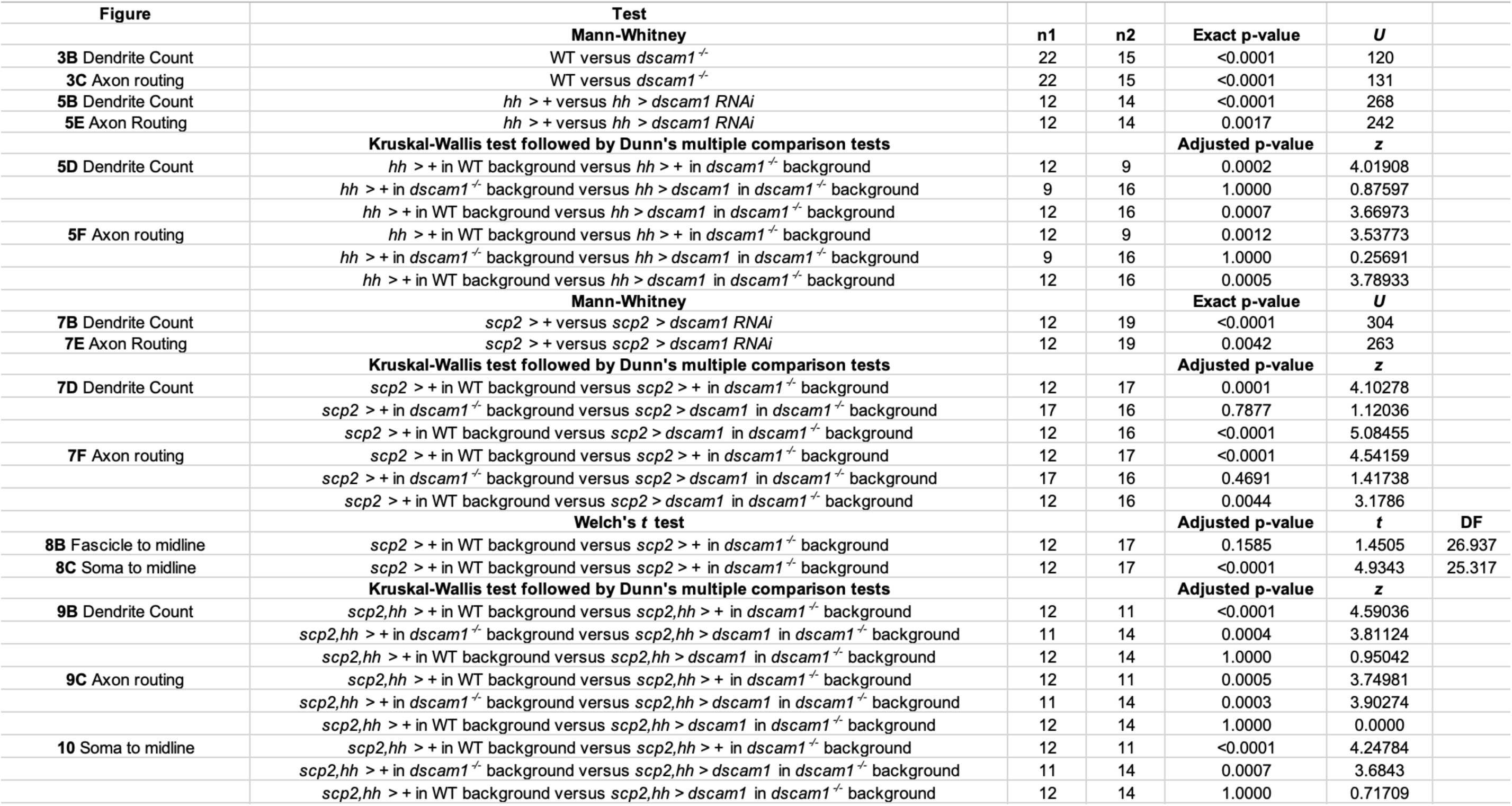
Statistical analyses grouped by figure number and panel, and statistical tests.

## Acknowledgments

We thank K. Banzai, M. Inal, O. Avraham, and the members of the Kamiyama lab for their insightful comments on the manuscript, and M. Fitch for technical support. This work was funded by the NIH grant (R01NS107558) to DK.

## Author contributions

KCB: Investigation, Methodology, Data curation, Visualization; Formal analysis, Writing – original draft, Writing – review and editing. DK: Writing – review and editing, Funding acquisition, Supervision.

## Competing interests

The authors declare no competing financial interests.

## Notes

### Competing Interest Statement

The authors have declared no competing interest.

## References

Alavi M, Song M, King GL, Gillis T, Propst R, Lamanuzzi M, Bousum A, Miller A, Allen R, Kidd T (2016) Dscam1 Forms a Complex with Robo1 and the N-Terminal Fragment of Slit to Promote the Growth of Longitudinal Axons. PLoS Biol 14:e1002560.

Andrews GL, Tanglao S, Farmer WT, Morin S, Brotman S, Berberoglu MA, Price H, Fernandez GC, Mastick GS, Charron F, Kidd T (2008) Dscam guides embryonic axons by Netrin-dependent and -independent functions. Development 135:3839–3848.

Avraham O, Hadas Y, Vald L, Hong S, Song MR, Klar A (2010) Motor and dorsal root ganglion axons serve as choice points for the ipsilateral turning of dI3 axons. J Neurosci 30:15546–15557.

Avraham O, Hadas Y, Vald L, Zisman S, Schejter A, Visel A, Klar A (2009) Transcriptional control of axonal guidance and sorting in dorsal interneurons by the Lim-HD proteins Lhx9 and Lhx1. Neural Dev 4:21.

Brose K, Bland KS, Wang KH, Arnott D, Henzel W, Goodman CS, Tessier-Lavigne M, Kidd T (1999) Slit proteins bind Robo receptors and have an evolutionarily conserved role in repulsive axon guidance. Cell 96:795–806.

Cohen O, Vald L, Yamagata M, Sanes JR, Klar A (2017) Roles of DSCAM in axonal decussation and fasciculation of chick spinal interneurons. Int J Dev Biol 61:235–244.

Constance WD, Mukherjee A, Fisher YE, Pop S, Blanc E, Toyama Y, Williams DW (2018) Neurexin and Neuroligin-based adhesion complexes drive axonal arborisation growth independent of synaptic activity. Elife 7.

Dascenco D, Erfurth ML, Izadifar A, Song M, Sachse S, Bortnick R, Urwyler O, Petrovic M, Ayaz D, He H, Kise Y, Thomas F, Kidd T, Schmucker D (2015) Slit and Receptor Tyrosine Phosphatase 69D Confer Spatial Specificity to Axon Branching via Dscam1. Cell 162:1140–1154.

Dong H, Guo P, Zhang J, Wu L, Fu Y, Li L, Zhu Y, Du Y, Shi J, Zhang S, Li G, Xu B, Bian L, Zhu X, You W, Shi F, Yang X, Huang J, Jin Y (2022) Self-avoidance alone does not explain the function of Dscam1 in mushroom body axonal wiring. Curr Biol 32:2908–2920 e2904.

Evans TA (2016) Embryonic axon guidance: insights from Drosophila and other insects. Curr Opin Insect Sci 18:11–16.

Evans TA, Bashaw GJ (2010) Axon guidance at the midline: of mice and flies. Curr Opin Neurobiol 20:79–85.

Fazeli A, Dickinson SL, Hermiston ML, Tighe RV, Steen RG, Small CG, Stoeckli ET, Keino-Masu K, Masu M, Rayburn H, Simons J, Bronson RT, Gordon JI, Tessier-Lavigne M, Weinberg RA (1997) Phenotype of mice lacking functional Deleted in colorectal cancer (Dcc) gene. Nature 386:796–804.

Furrer MP, Kim S, Wolf B, Chiba A (2003) Robo and Frazzled/DCC mediate dendritic guidance at the CNS midline. Nat Neurosci 6:223–230.

Furrer MP, Vasenkova I, Kamiyama D, Rosado Y, Chiba A (2007) Slit and Robo control the development of dendrites in Drosophila CNS. Development 134:3795–3804.

Goyal G, Zierau A, Lattemann M, Bergkirchner B, Javorski D, Kaur R, Hummel T (2019) Inter-axonal recognition organizes Drosophila olfactory map formation. Sci Rep 9:11554.

Guan KL, Rao Y (2003) Signalling mechanisms mediating neuronal responses to guidance cues. Nat Rev Neurosci 4:941–956.

Harris R, Sabatelli LM, Seeger MA (1996) Guidance cues at the Drosophila CNS midline: identification and characterization of two Drosophila Netrin/UNC-6 homologs. Neuron 17:217–228.

Hattori D, Demir E, Kim HW, Viragh E, Zipursky SL, Dickson BJ (2007) Dscam diversity is essential for neuronal wiring and self-recognition. Nature 449:223–227.

Hattori D, Chen Y, Matthews BJ, Salwinski L, Sabatti C, Grueber WB, Zipursky SL (2009) Robust discrimination between self and non-self neurites requires thousands of Dscam1 isoforms. Nature 461:644–648.

Haverkamp LJ (1986) Anatomical and physiological development of the Xenopus embryonic motor system in the absence of neural activity. J Neurosci 6:1338–1348.

Hiramoto M, Hiromi Y, Giniger E, Hotta Y (2000) The Drosophila Netrin receptor Frazzled guides axons by controlling Netrin distribution. Nature 406:886–889.

Howard LJ, Brown HE, Wadsworth BC, Evans TA (2019) Midline axon guidance in the Drosophila embryonic central nervous system. Semin Cell Dev Biol 85:13–25.

Inal MA, Banzai K, Kamiyama D (2020) Retrograde Tracing of Drosophila Embryonic Motor Neurons Using Lipophilic Fluorescent Dyes. J Vis Exp.

Inal MA, Bui KC, Marar A, Li S, Kner P, Kamiyama D (2021) Imaging of In Vitro and In Vivo Neurons in Drosophila Using Stochastic Optical Reconstruction Microscopy. Curr Protoc 1:e203.

Kamiyama D, McGorty R, Kamiyama R, Kim MD, Chiba A, Huang B (2015) Specification of Dendritogenesis Site in Drosophila aCC Motoneuron by Membrane Enrichment of Pak1 through Dscam1. Dev Cell 35:93–106.

Kaprielian Z, Runko E, Imondi R (2001) Axon guidance at the midline choice point. Dev Dyn 221:154–181.

Kidd T, Bland KS, Goodman CS (1999) Slit is the midline repellent for the robo receptor in Drosophila. Cell 96:785–794.

Kidd T, Brose K, Mitchell KJ, Fetter RD, Tessier-Lavigne M, Goodman CS, Tear G (1998) Roundabout controls axon crossing of the CNS midline and defines a novel subfamily of evolutionarily conserved guidance receptors. Cell 92:205–215.

Klambt C, Jacobs JR, Goodman CS (1991) The midline of the Drosophila central nervous system: a model for the genetic analysis of cell fate, cell migration, and growth cone guidance. Cell 64:801–815.

Kolodziej PA, Timpe LC, Mitchell KJ, Fried SR, Goodman CS, Jan LY, Jan YN (1996) frazzled encodes a Drosophila member of the DCC immunoglobulin subfamily and is required for CNS and motor axon guidance. Cell 87:197–204.

Landgraf M, Bossing T, Technau GM, Bate M (1997) The origin, location, and projections of the embryonic abdominal motorneurons of Drosophila. J Neurosci 17:9642–9655.

Landgraf M, Sanchez-Soriano N, Technau GM, Urban J, Prokop A (2003a) Charting the Drosophila neuropile: a strategy for the standardised characterisation of genetically amenable neurites. Dev Biol 260:207–225.

Landgraf M, Jeffrey V, Fujioka M, Jaynes JB, Bate M (2003b) Embryonic origins of a motor system: motor dendrites form a myotopic map in Drosophila. PLoS Biol 1:E41.

Lanoue V, Cooper HM (2019) Branching mechanisms shaping dendrite architecture. Dev Biol 451:16–24.

Lefebvre JL, Sanes JR, Kay JN (2015) Development of dendritic form and function. Annu Rev Cell Dev Biol 31:741–777.

Liu C, Trush O, Han X, Wang M, Takayama R, Yasugi T, Hayashi T, Sato M (2020) Dscam1 establishes the columnar units through lineage-dependent repulsion between sister neurons in the fly brain. Nat Commun 11:4067.

Liu G, Li W, Wang L, Kar A, Guan KL, Rao Y, Wu JY (2009) DSCAM functions as a netrin receptor in commissural axon pathfinding. Proc Natl Acad Sci U S A 106:2951–2956.

Long KR, Huttner WB (2019) How the extracellular matrix shapes neural development. Open Biol 9:180216.

Mauss A, Tripodi M, Evers JF, Landgraf M (2009) Midline signalling systems direct the formation of a neural map by dendritic targeting in the Drosophila motor system. PLoS Biol 7:e1000200.

Mitchell KJ, Doyle JL, Serafini T, Kennedy TE, Tessier-Lavigne M, Goodman CS, Dickson BJ (1996) Genetic analysis of Netrin genes in Drosophila: Netrins guide CNS commissural axons and peripheral motor axons. Neuron 17:203–215.

Organisti C, Hein I, Grunwald Kadow IC, Suzuki T (2015) Flamingo, a seven-pass transmembrane cadherin, cooperates with Netrin/Frazzled in Drosophila midline guidance. Genes Cells 20:50–67.

Saalfeld S, Cardona A, Hartenstein V, Tomancak P (2009) CATMAID: collaborative annotation toolkit for massive amounts of image data. Bioinformatics 25:1984–1986.

Sanes JR (1989) Extracellular matrix molecules that influence neural development. Annu Rev Neurosci 12:491–516.

Schmucker D, Clemens JC, Shu H, Worby CA, Xiao J, Muda M, Dixon JE, Zipursky SL (2000) Drosophila Dscam is an axon guidance receptor exhibiting extraordinary molecular diversity. Cell 101:671–684.

Sink H, Whitington PM (1991) Location and connectivity of abdominal motoneurons in the embryo and larva of Drosophila melanogaster. J Neurobiol 22:298–311.

Sun W, You X, Gogol-Doring A, He H, Kise Y, Sohn M, Chen T, Klebes A, Schmucker D, Chen W (2013) Ultra-deep profiling of alternatively spliced Drosophila Dscam isoforms by circularization-assisted multi-segment sequencing. EMBO J 32:2029–2038.

Sundararajan L, Stern J, Miller DM, 3rd (2019) Mechanisms that regulate morphogenesis of a highly branched neuron in C. elegans. Dev Biol 451:53–67.

Tomer R, Khairy K, Amat F, Keller PJ (2012) Quantitative high-speed imaging of entire developing embryos with simultaneous multiview light-sheet microscopy. Nat Methods 9:755–763.

Varoqueaux F, Sigler A, Rhee JS, Brose N, Enk C, Reim K, Rosenmund C (2002) Total arrest of spontaneous and evoked synaptic transmission but normal synaptogenesis in the absence of Munc13-mediated vesicle priming. Proc Natl Acad Sci U S A 99:9037–9042.

Verhage M, Maia AS, Plomp JJ, Brussaard AB, Heeroma JH, Vermeer H, Toonen RF, Hammer RE, van den Berg TK, Missler M, Geuze HJ, Sudhof TC (2000) Synaptic assembly of the brain in the absence of neurotransmitter secretion. Science 287:864–869.

Wilhelm N, Kumari S, Krick N, Rickert C, Duch C (2022) Dscam1 Has Diverse Neuron Type-Specific Functions in the Developing Drosophila CNS. eNeuro 9.

Winding M, Pedigo BD, Barnes CL, Patsolic HG, Park Y, Kazimiers T, Fushiki A, Andrade IV, Khandelwal A, Valdes-Aleman J, Li F, Randel N, Barsotti E, Correia A, Fetter RD, Hartenstein V, Priebe CE, Vogelstein JT, Cardona A, Zlatic M (2023) The connectome of an insect brain. Science 379:eadd9330.

Wojtowicz WM, Flanagan JJ, Millard SS, Zipursky SL, Clemens JC (2004) Alternative splicing of Drosophila Dscam generates axon guidance receptors that exhibit isoform-specific homophilic binding. Cell 118:619–633.

Wojtowicz WM, Wu W, Andre I, Qian B, Baker D, Zipursky SL (2007) A vast repertoire of Dscam binding specificities arises from modular interactions of variable Ig domains. Cell 130:1134–1145.

Yamagata M, Sanes JR (2008) Dscam and Sidekick proteins direct lamina-specific synaptic connections in vertebrate retina. Nature 451:465–469.

Yogev S, Shen K (2014) Cellular and molecular mechanisms of synaptic specificity. Annu Rev Cell Dev Biol 30:417–437.

Yogev S, Shen K (2017) Establishing Neuronal Polarity with Environmental and Intrinsic Mechanisms. Neuron 96:638–650.

Zhan XL, Clemens JC, Neves G, Hattori D, Flanagan JJ, Hummel T, Vasconcelos ML, Chess A, Zipursky SL (2004) Analysis of Dscam diversity in regulating axon guidance in Drosophila mushroom bodies. Neuron 43:673–686.

Zhu H, Hummel T, Clemens JC, Berdnik D, Zipursky SL, Luo L (2006) Dendritic patterning by Dscam and synaptic partner matching in the Drosophila antennal lobe. Nat Neurosci 9:349–355.

Zou W, Shen A, Dong X, Tugizova M, Xiang YK, Shen K (2016) A multi-protein receptor-ligand complex underlies combinatorial dendrite guidance choices in C. elegans. Elife 5.

